# Multiomic Data Integration Reveals Microbial Drivers of Aetiopathogenesis in Mycosis Fungoides

**DOI:** 10.1101/2024.01.15.575621

**Authors:** Philipp Licht, Volker Mailänder

## Abstract

**Background:** Mycosis fungoides (MF) represents the most prevalent entity of cutaneous T cell lymphoma (CTCL). The MF aetiopathogenesis is incompletely understood, due to strong transcriptomic heterogeneity and opposing perspectives on the initial oncologic transformation mapping the event to both early thymocytes and mature, effector memory T cells. Recently, using clinical specimen, our group showed that the skin microbiome aggravates disease course, mainly driven by an outgrowing, pathogenic *S. aureus* strain carrying the virulence factor spa, which reportedly activates the T cell signalling pathway NF-κB.

**Methods:** To further investigate the role of the skin microbiome in MF aetiopathogenesis, we here performed RNA sequencing, multiomic data integration of the skin microbiome and skin transcriptome using Multi-Omic Factor Analysis (MOFA), virome profiling, and T cell receptor (TCR) sequencing in 10 MF patients representing a subset of our previous study cohort.

**Results:** We observed that inter-patient transcriptional heterogeneity may be largely driven by differential activation of T cell signalling pathways. Strikingly, the MOFA model resolved the heterogenous activation pattern of T cell signalling after denoising the transcriptome from microbial influence. The MOFA model showed that the outgrowing *S. aureus* strain evoked signalling by non-canonical NF-κB and IL-1B, which likely fuelled the aggravated disease course. Further, the MOFA model revealed aberrant pathways of early thymopoiesis alongside enrichment of antiviral innate immunity. In line, viral prevalence, particularly of Epstein-Barr virus (EBV), trended higher in both lesional skin and the blood compared to nonlesional skin. Additionally, TCRs in both MF skin lesions and the blood were significantly more likely to recognize EBV peptides involved in latent infection.

**Conclusions:** First, our findings suggest that *S. aureus* with its virulence factor spa fuels MF progression trough non-canonical NF-κB and IL-1B signalling. Second, our data provide insights into the potential role of viruses in MF aetiology. Last, we propose a model of microbiome-driven MF aetiopathogenesis: Thymocytes undergo initial oncologic transformation, potentially caused by viruses. After maturation and skin infiltration, an outgrowing, pathogenic *S. aureus* strain evokes activation and maturation into effector memory T cells, resulting in aggressive disease.

## Introduction

Cutaneous T cell lymphoma (CTCL) is a heterogeneous group of non-Hodgkin T cell lymphomas with skin homing properties. The most common entity is Mycosis fungoides (MF) with an incidence rate of 4.1 cases per million in the USA (*1*, *2*). MF patients present with several to many cutaneous lesions that are formed by the infiltration of neoplastic T cells and benign reactive lymphocytes. With disease progression, both neoplastic and benign infiltrate accumulate, resulting in inflammatory reddening of skin lesions (*1*, *3*, *4*). Depending on the degree of lymphocyte infiltration and inflammation, lesions are classified into the stages patch, plaque, and tumour. As MF is a lymphoproliferative disorder which can involve extracutaneous compartments like the blood, a clinical staging system considers these events and classifies patients into stages IA – IVB (*1*, *5*). In early stages (IA – IIA), MF is an indolent disease with a 5-year disease-specific survival of 89%, which, however, dramatically drops to ∼20% in the most advanced stages (*5*). Because the aetiology and pathogenesis of MF are incompletely understood, treatment options are limited, and cure is almost not achievable (*6*, *7*).

MF is thought to arise from mature, skin-resident CD4+ T cells (*8*), which resemble the phenotype of tissue resident effector memory T cells (*9*). However, others suggest that the initial oncologic transformation takes place during early thymopoiesis, specifically during the stages double-negative (DN) DN-1 through DN-3. Those “premalignant clones” may then migrate to the skin where they proliferate (*10–15*). The causative agent responsible for the oncogenic transformation of T cells in MF remains uncertain (*1*). Viruses are among the potential factors considered due to their involvement in various lymphoma types, including even two other entities of CTCL (*16*). Viral involvement in MF is further supported by an elevated risk of MF patients to develop one or more virus-initiated lymphoma types either simultaneously with MF or later in time (*17–23*). However, the role of viruses in the aetiology of MF remains unclear, as findings from different studies have yielded conflicting results (*24–26*).

Likewise, the molecular drivers of MF pathogenesis remain incompletely understood (*27*, *28*). In normal T cells, three signals orchestrate activity and proliferation: First, an initial T cell response is initiated by antigenic stimulation of the T cell receptor (TCR) together with CD3. Second, co-stimulation is required to augment TCR signalling, which is mediated by various molecules, including CD28 and the tumour necrosis factor receptor super family (TNFRSF). Downstream, both TCR signalling and co-stimulation converge on PI3K/AKT, NFAT and NF-κB (*29*). Third, sustained T cell activity is promoted by cytokines, which activate JAK-STAT. While CD8+ T cells require interleukin (IL) 12 and interferon (IFN) α/β to initiate signal three, CD4+ T cells require IL-1 (*30*, *31*). Because these pathways are frequently dysregulated in T cell lymphomas, a “three-signal model” of T cell lymphoma pathogenesis has been proposed (*32*). In MF, dysregulation of TCR, TNFRSF/NF-κB and JAK-STAT pathways are recurrently observed, demonstrating the involvement of all three “T cell lymphoma promoting” signalling pathways (*33–38*). However, the MF transcriptome exhibits substantial variability between patients and among lesions within the same patient (*27*, *39*), which impedes the identification of a shared pathogenic pathway to date.

It has been proposed that the skin microbiome contributes to or evokes transcriptional heterogeneity (*40*). In agreement, we recently identified a subgroup of MF patients with a significantly aggravated disease course and outgrowth of a distinct, pathogenic *S. aureus* strain on plaque lesions. Conversely, another MF patient subgroup presented with a more balanced skin microbiome and a favourable prognosis. Reflecting the differing prevalences of *S. aureus* between the two subgroups, we referred to the subgroup with aggravated disease and *S. aureus* outgrowth as ΔSA-positive, while the other subgroup was termed ΔSA-neutral. Notably, the virulence factor staphylococcal protein A (spa) was highly abundant in the genome of *S. aureus* outgrowing in the ΔSA-positive subgroup (*41*). It has been demonstrated that spa activates the NF-κB pathway (*42*, *43*), which is involved in T cell co-stimulation (*29*), a component of the “three-signal model” of T cell lymphoma pathogenesis (*32*). Moreover, some studies observed aberrant NF-κB activity in subsets of MF patients with aggressive disease (*44–46*). We thus theorized that the skin microbiome shapes MF disease signalling, resulting in exacerbated malignancy.

To investigate our hypothesis, we here performed bulk RNA sequencing (RNAseq), multiomic data integration of the microbiome and the transcriptome using Multi-Omic Factor Analysis (MOFA) (*48*), virome profiling and T cell receptor sequencing (TCRseq). Our analyses yielded three main findings.

First, our data indicated that inter-patient transcriptional heterogeneity was largely driven by differential expression of pathways involved in T cell signalling. Strikingly, denoising the transcriptome from microbial influence using MOFA pronouncedly reduced the heterogeneous activation pattern of T cell signalling pathways. This strongly suggested that the skin microbiome had a substantial impact on MF disease signalling.

Second, the MOFA model suggested that *S. aureus* with its virulence factor spa induced ectopic activity of both non-canonical NF-κB and IL-1B signalling. While non-canonical NF-κB signalling leads to survival, proliferation and differentiation of naïve T cell into effector memory T cells (*49*), IL-1B facilitates sustained CD4+ T cell activation (*30*, *31*). Given that CD4+ effector memory T cells are the malignant T cell subset in MF (*9*), spa-bearing *S. aureus* may induce or augment the phenotypic characteristics of MF, resulting in aggressive disease.

Third, the MOFA model uncovered active anti-viral immune response along with enriched pathways involved in early thymopoiesis between DN1 and DN3. Viral prevalence, particularly Epstein-Barr-Virus (EBV) and Human Papillomavirus 71 (HPV), trended higher in both lesional skin and the blood. In line, TCRs in both MF skin lesions and blood were significantly more likely to recognize epitopes of EBV compared to TCRs in nonlesional skin. Notably, the most frequently recognized EBV-epitopes were derived from proteins orchestrating latent EBV-infection. Our findings collectively provide evidence supporting the potential viral involvement in the aetiology of MF, considering that (I) malignant MF T cells infiltrate the skin from the blood (*50*), (II) the initial oncologic transformation of malignant T cells in MF was mapped between DN1 to DN3 (*10–15*), and (III) latent EBV infection can induce non-Hodgkin lymphoma, including peripheral T cell lymphoma (*47*, *48*).

Last, we propose a model of microbiome-driven MF aetiopathogenesis: The initial oncologic transformation occurs during early thymopoiesis, possibly induced by viral infection. Following maturation and skin infiltration, outgrowth of a spa-bearing *S. aureus* strain exacerbates disease by activating both non-canonical NF-κB and IL-1B signalling, resulting in the differentiation of naïve T cells into effector memory T cells with sustained activity. Our study highlights the critical role of the microbiome in MF aetiopathogenesis.

## Results

### Study cohort and clinical specimens

The study cohort consisted of 10 MF patients (mean age at sampling: 64.7 years; range: 47 to 82 years) and was a subset of patients that were enrolled in our previous investigation where we characterized the MF skin microbiome (*41*). To ensure traceability, IDs of included patients in this study were adopted from our previous investigation (*41*). In that study, we found that a subgroup of patients presented with an increased abundance of *S. aureus* on MF lesions compared to nonlesional skin of the same patient. We termed this subgroup ΔSA-positive as opposed to the ΔSA-neutral subgroup, where *S. aureus* abundance was consistent between MF lesions and nonlesional skin of the same patient. The ΔSA-positive subgroup had a strong dysbiosis and a significantly aggravated disease course compared to the ΔSA-neutral subgroup (*41*). Of the 10 MF patients included in the present study, three were in the ΔSA-positive subgroup and seven were in the ΔSA-neutral subgroup.

Clinical specimens included skin swabs and punch biopsies of lesional skin (five patch stage and five plaque stage) and matched nonlesional skin of the same MF patients as intra-patient control (10 nonlesional skin samples). For more details about the rational of using internal controls, please refer to Licht, Dominelli et al. (*41*). Additionally, blood was drawn in four patients for the isolation of peripheral blood mononuclear cells (PBMCs). All clinical specimens were taken at the same visit. Skin punch biopsies were used for RNAseq. Where sufficient material was available, skin punch biopsies and PBMCs were used for TCRseq. Shotgun metagenomics from skin swabs were initially carried out in our previous investigation (*41*) and re-analysed in the present study in a multi-omic data integration approach together with the skin transcriptome. Table 1 summarizes patient metadata, clinical specimens, and data modalities used in the present study.

**Table 1:** Study cohort, clinical specimens, and patient metadata. This study cohort is a subset of the cohort enrolled in Licht et al. (*41*), in which we characterized the skin microbiome of MF patients. We demonstrated that *S. aureus* abundance is increases on MF lesions compared to nonlesional skin in a subgroup of patients (ΔSA-positive) while *S. aureus* abundance does not change in the other subgroup (ΔSA-neutral). ΔSA-positive patients exhibit a poor clinical course compared to ΔSA-neutral patients. Patient IDs in this study match the given patient IDs in Licht et al. (*41*). NA = not available; f = female; m = males

| Patient ID | Sex | Age (years) | Clinical Stage | ΔSA-subgroup | Stage of lesion (Body Site) | Blood Sample | TCRseq: Tissues included |
| --- | --- | --- | --- | --- | --- | --- | --- |
| Pat1 | f | 81 | IA | positive | Plaque (hip) | Yes | Skin (Plaque), Blood |
| Pat3 | f | 47 | IB | neutral | Patch (thigh) | No | NA |
| Pat4 | m | 70 | IIB | positive | Plaque (upper back) | Yes | Skin (Plaque and nonlesional), Blood |
| Pat5 | f | 73 | IB | neutral | Patch (thigh) | No | Skin (Patch and nonlesional) |
| Pat7 | m | 54 | IB | neutral | Plaque (flank) | Yes | Skin (Plaque and nonlesional) |
| Pat8 | m | 82 | IIB | positive | Patch (upper back) | Yes | Skin (Patch), Blood |
| Pat9 | m | 63 | IB | neutral | Plaque (forearm) | No | Skin (Plaque) |
| Pat10 | m | 49 | IB | neutral | Patch (gluteus) | No | Skin (Patch) |
| Pat13 | m | 66 | IB | neutral | Patch (lower leg, front) | No | NA |
| Pat14 | m | 62 | IB | neutral | Plaque (lower leg, front) | No | Skin (Plaque and nonlesional), Blood |

### Transcriptional heterogeneity may be largely driven by differential activation of pathways of the “three-signal model” of T cell lymphoma pathogenesis

A number of studies investigated the CTCL transcriptome (*27*, *39*) and uncovered various mechanisms of disease progression (*33–38*). However, the transcriptome was also found to exhibit significant heterogeneity among patients, complicating the identification of a common pathological mechanism (*27*, *39*, *46*, *49–53*).

To explore the root of transcriptional heterogeneity, we employed RNAseq together with single gene analysis and gene set enrichment analysis (GSEA). Principal component analysis (PCA) of the MF skin transcriptome clearly separated nonlesional skin, patch, and plaque in principal component (PC) 1. However, lesional samples also spread along PC2, demonstrating strong inter-patient transcriptional heterogeneity (Fig. 1A). Likewise, GSEA recovered several aberrant pathways known to promote MF exacerbation, such as chemokine signalling (*54*) (Suppl. Material 2), but also showed substantial activation differences between patients in both patch and plaque lesions (Suppl. Material 3). Notably, pathways of the “three-signal model” of T cell lymphoma pathogenesis exhibited different activation patterns between patch and plaque (Fig. 1B): TCR signalling and co-stimulation by CD28 were enriched in both patch and plaque, whereas Interleukin 13 (IL-13) signalling and co-stimulation through death receptors activating NF-κB (*44, 45*) were enriched solely in plaque stage.

**Figure 1:**
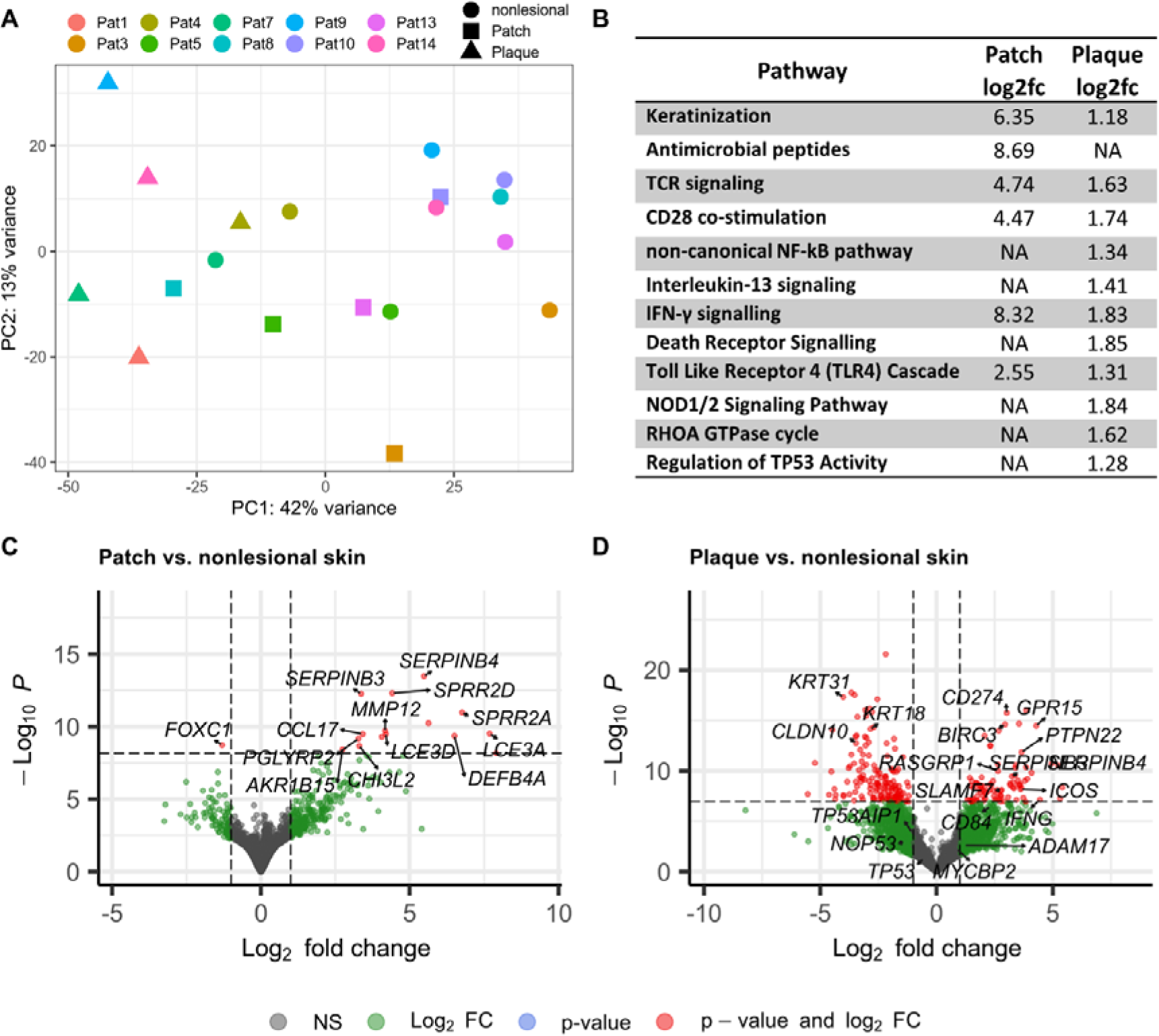
Analysis of the transcriptome. **A)** Principal component analysis showing inter-heterogeneity of plaque stage **B)** Gene set enrichment analysis **C-D)** Volcano plots showing the distribution of differentially expressed genes. PC = Principal Component, Log2fc = log2 fold-change, NA = not available

Next, we aimed to assess whether pathways of the “three-signal model” for T cell lymphoma pathogenesis (*32*) replicate transcriptional heterogeneity. To this end, we compiled a list of genes that are (I) members of the “three-signal model” for T cell lymphomas and (II) were differentially expressed in either or both patch and plaque stage. A clustered heatmap of these genes closely resembled the pattern of the PCA of the entire transcriptome: Nonlesional skin and plaque samples formed separate clusters, while patch samples exhibited intermediate expression levels. Importantly, plaque samples displayed pronounced heterogeneity in RNA expression levels (Fig. 2A). We thus reasoned that inter-patient transcriptional heterogeneity can, at least to some extent, be attributed to differential expression of pathways that orchestrate T cell lymphoma pathogenesis.

**Figure 2:**
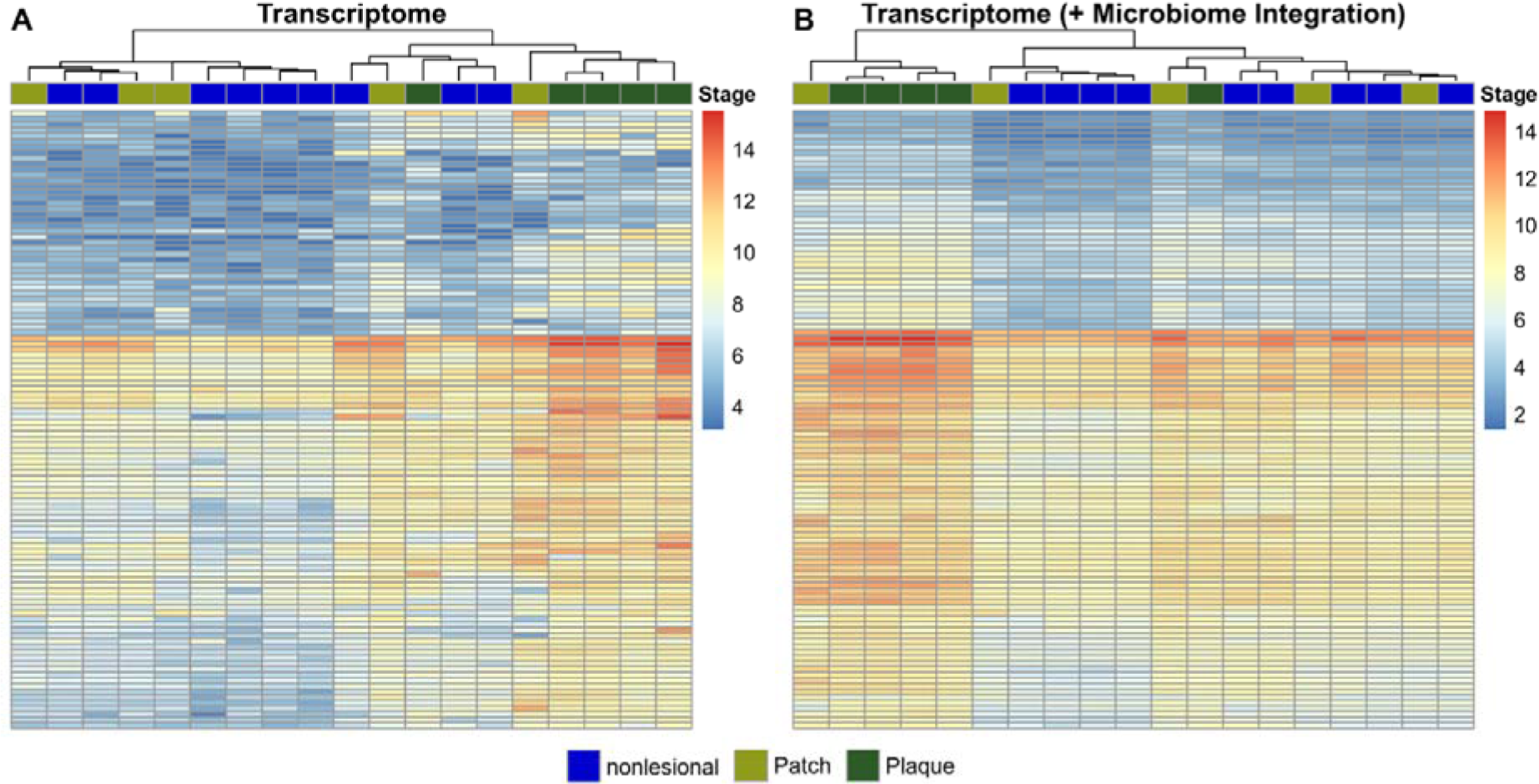
Clustered heatmap of genes involved in the “three-signal model” of T cell lymphoma pathogenesis. The clustered heatmap was created with genes involved in the “three-signal model” of T cell lymphoma pathogenesis that were differentially expressed in patch and/or plaque stage and were included in the MOFA model (5000 highly variable genes, see methods and suppl. Material 6). **A)** Shown are normalized RNA expression levels before integration of the microbiome. Notably, although only genes of the “three-signal model” of T cell lymphoma pathogenesis were included, the clustering resembled the patterns of the PCA, which was based on the entire transcriptome. Despite clustering together, plaque samples exhibited a high degree of heterogeneity. **B)** After data integration of the transcriptome with the microbiome, the heterogeneity of plaque samples was largely resolved suggesting that the microbiome had a strong impact on MF disease signalling.

### The MF skin transcriptome shows responses to microbial stimuli

Since differential microbial skin colonization may contribute to transcriptional heterogeneity (*40*), and we recently identified that the skin microbiome stratifies MF patients and determines the clinical course (*41*), our next objective was to identify microbial responses in the skin transcriptome.

As expected, we uncovered enriched pathways likely activated by the skin microbiome (Fig. 1B and suppl. Material 2). Notably, Toll Like Receptor 4 (TLR4) was upregulated in patch and showed attenuated activity in plaque. The pathway senses lipopolysaccharides, a cell wall constituent of gram-negative bacteria, and activates activates inflammatory responses as well as pyroptosis, which is a form of programmed necrosis. Pyroptosis protects the host from microbial infection but, if overactivated, can also lead to pathological inflammation (*55*), a typical condition in progressive MF (*4*). In accordance, we recently showed that the skin microbiome of patches is strongly dysbiotic, while dysbiosis on plaques was observed only on a subgroup of MF patients (*41*), providing an explanation for the differential TLR4 activation between patch and plaque.

Conversely, several pathways were solely upregulated in plaque, likely activated by the distinct, *S. aureus* strain carrying the virulence factors α-hemolysin and spa, which outgrows on plaque of the ΔSA-positive subgroup (*41*). NOD2 (nucleotide-binding oligomerization domain 2) senses small peptides of the cell wall component peptidoglycan found in gram-positive bacteria such as *S. aureus*. Notably, protection specifically against this pathogen is facilitated by the interaction of NOD2 with the *S. aureus* virulence factor α-hemolysin (*56*, *57*). Further, we observed increased activity of the RHOA GTPase cycle, which can be hijacked by *S. aureus* through its virulence factor spa to invade epithelial barriers (*58*). Additionally, we detected an enrichment of IL-13 signalling, which was shown to be secreted by the skin following exposure to *S. aureus* (*59*, *60*). In agreement, it was reported that malignant T cells in the skin of MF patients expressed IL-13, whereas malignant T cells in the lymph nodes and blood did not (*46*).

Collectively, the transcriptome revealed responses to microbial stimuli that aligned with the microbiome patterns identified on MF lesions in our prior study (*41*). Consequently, our findings indicated that distinct microbial colonization led to differential transcriptomic responses between patients.

### Multi-omic data integration of the microbiome and the transcriptome resolves transcriptional heterogeneity

To investigate whether differential skin colonization elicited the heterogeneous expression of genes related to the “three-signal model” for T cell lymphoma pathogenesis, we performed data integration of the microbiome and the transcriptome using Multi-Omic Factor Analysis (MOFA) (*61*). Briefly, MOFA can be seen as a multi-omic implementation of PCA and finds latent factors (comparable to principal components in PCA) that capture the main sources of variation across different omic data modalities (which are called views in the MOFA framework). Within the latent factors, weights are allocated to the features of the different views. Consequently, latent factors are characterized by feature weights, which indicate their importance or significance in the variation captured by the latent factor. Downstream, additional analyses such as gene set enrichment analysis (GSEA) can be conducted within each latent factor. The MOFA framework further enables the identification of each view’s contribution to the variation captured by a latent factor across the entire multi-omic dataset (*61*).

Regarding the multi-omic data set in this study, the weights of the microbes (features of the microbiome) and the weights of the genes (features of the transcriptome) characterize the latent factors. The MOFA model showed a good fit to the multi-omic data set, as the patterns of the latent factors were congruent to the results observed in both the independent transcriptome analysis (see above) and our previous study on the MF skin microbiome (*41*) (see Suppl. Material 4). Latent factors 1 to 5 captured shared sources of variation present in both data modalities, indicating a reciprocal influence between the transcriptome and microbiome (Fig. 3A). By leveraging the latent factors, MOFA reconstructs the input data to separate shared and specific variations in each data modality, thereby denoising the data and revealing underlying biological signals (*61*). Strikingly, the differential expression of genes involved in the “three-signal model” of T cell lymphoma pathogenesis was resolved after integration of the microbiome and transcriptome data modalities (Fig. 2B). This demonstrated that the lesion-specific microbiome heavily influences MF disease signalling.

**Figure 3:**
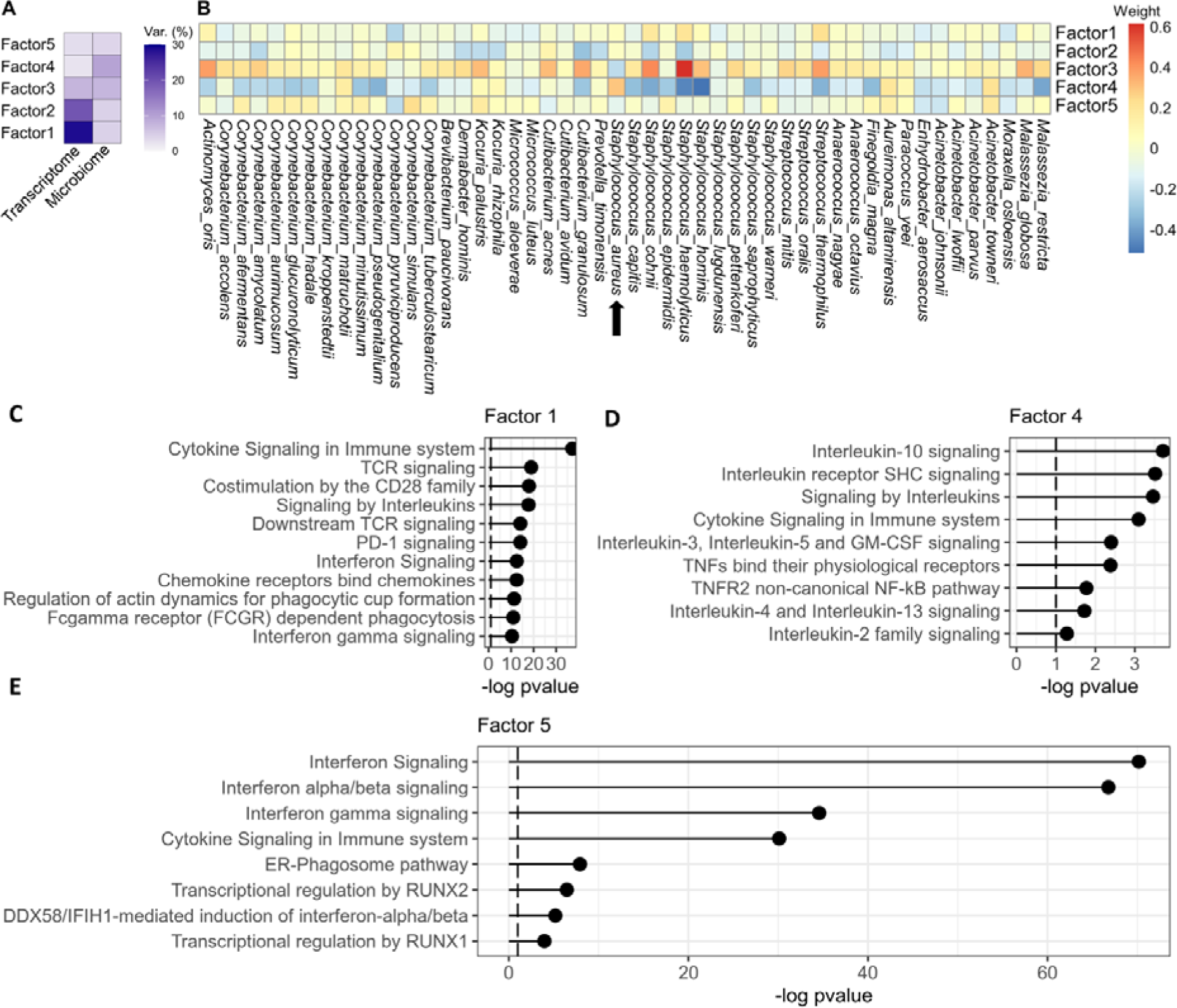
Overview of the latent factors of the MOFA model in the microbiome and transcriptome data modalities. **A)** Overview of the latent factors capturing variances in the microbiome and transcriptome data modalities. Factor 4 captured the majority of variance in the microbiome, meaning that this factor captured important impacts of the microbiome on the transcriptome. **B)** Overview of the feature weights of the microbiome per factor. **C – E)** Gene set enrichment analysis. Factor 1 showed pathomechanisms of high and sustained T cell activity in MF (TCR signalling, CD28 co-stimulation, interleukin signals). Factor 4 showed pathways that were likely evoked by *S. aureus*. Factor 5 showed strong anti-virus responses along with enrichment of RUNX1 and RUNX2 signalling, which have pivotal roles in thymopoiesis (*62–65*).

### Spa-bearing *S. aureus* activates non-canonical NF-**κ**B and IL-1B signalling

We next assessed the impact of the microbiome on MF disease signalling in more detail. **Factor 4** accounted for a substantial proportion of the total variance in the microbiome (Fig. 3A), and *S. aureus* was the only microbiome feature with a positive weight. Notably, microbes with anti-*S. aureus* properties, such as *S. hominis* and *S. epidermidis* (*66–68*), displayed markedly negative weights. Thus, the microbiome feature weights exhibited a pattern similar to plaque lesions of the ΔSA-positive subgroup. These patients presented with *S. aureus* outgrowth and an aggravated clinical course (*41*).

In the transcriptome, GSEA showed enrichment of both interleukin signalling, particularly IL-1B, and non-canonical NF-κB signalling (Fig. 3D, Suppl. Material 5), which control T cell activity. While non-canonical NF-κB serves as co-stimulator and generates and maintains effector memory T cells (*69*, *70*), IL-1B signalling facilitates sustained activation of CD4+ T cells (*32*). In agreement, the malignant T cell subset in MF is thought to be CD4+ effector memory T cells (*9*). Regarding IL-1B signalling, Chng et al. (*71*) reported that human keratinocytes express IL-1B when challenged with *S. aureus*. This suggests that outgrowth of virulent *S. aureus* stimulates the tumour microenvironment to activate malignant T cells via IL-1B in a paracrine fashion. Regarding non-canonical NF-κB signalling, we intriguingly observed that the stimulating ligand, TNF-α (*72*), displayed an almost neutral weight in factor 4 (Suppl. Material 5), indicating an alternative stimulation route. Instead, we previously identified high abundance of the virulence factor staphylococcal protein A (spa) in the genome of the *S. aureus* strain colonizing ΔSA-positive patients. It was reported that spa can activate the NF-κB pathway (*42*, *43*), and some studies associated aggressive CTCL with upregulated NF-κB signalling (*44–46*). Notably, several genes emphasized in these studies exhibited positive weights in factor 4 (e.g., *LTA, LTB, BIRC3, TNFSF13, TNFSF14, TNFRSF1B* and *TNFRSF7*; Suppl. Material 5). Owing to the almost neutral weight of the NF-κB stimulating agent TNF-α and the presence of spa as an alternative stimulator (*41*), we theorized that outgrowing, spa-bearing *S. aureus* strains upregulated non-canonical NF-κB signalling in MF patients with aggressive disease.

Collectively, the MOFA model showed that the skin microbiome significantly influences MF disease signalling and is a major source of transcriptional heterogeneity. Further, the MOFA model suggested that *S. aureus* fuelled malignant T cells to promote the aggravated disease course of ΔSA-positive patients via two signalling axes: While spa activates non-canonical NF-κB signalling resulting in survival, proliferation, and naïve T cell differentiation into mature, effector memory T cells, the neoplastic T cell subset in MF (*9*), paracrine IL-1B signalling facilitates sustained activation.

### Aberrant signalling of early thymopoiesis alongside enriched anti-viral immunity suggests viral involvement in MF aetiology

Another interesting transcriptomic pattern was present in **factor 5.** While the microbiome weights showed only minimal signals (Fig. 3B), GSEA identified significant upregulation of host defence signals against viruses (Fig. 3E), indicating that factor 5 represented a source of variation independent of bacteria. In particular, GSEA showed upregulation of interferon alpha and beta (IFN-α/β) signalling, which is the first line of innate immune defence upon viral infection. Downstream, IFN-α/β signalling stimulates genes that inhibit the replication machinery of viruses at various mechanistic levels (*73*). Several of these interferon-stimulated genes exhibited high weights in factor 5, for example *MX1, IFIT1, IFIT3, OAS1, OAS2, IFI27,* and *OASL* (Suppl. Material 5). *IFIT3*, which displayed the highest weight in factor 5, was shown to specifically boost antiviral signalling by IFN-α/β (*74*). Further, GSEA found an enrichment of the ER-phagosome pathway (Fig. 3E), which can be hijacked by viruses for their own translation, replication and particle budding in order to spread into other host cells (*75*). Notably, the phagosome-pathway genes *TAP1* and *TAP2*, which displayed high weights in factor 5 (Suppl. Material 5), are highly expressed by Epstein-Barr-Virus (EBV) infected lymphocytes (*76*), and interact with Epstein-Barr-nuclear antigen 1 (EBNA1) (*77*). Interestingly, EBV infection can lead to *RUNX1* expression (*78*, *79*), which was significantly enriched along with *RUNX2* (Fig. 3E). The RUNX family is a frequent target of retroviral insertion (*80–82*), resulting in the development of several T cell lymphoma entities (*83*, *84*). *RUNX1* regulates the expansion of mature CD4+ T cells (*63*), which constitutes the neoplastic T cell subset in MF (*9*). Notably, both *RUNX1* and *RUNX2* are important regulators of early thymopoiesis during the double negative stages of T lymphocytes (*62–65*), a developmental step that has been mapped to the initial oncologic transformation of T cells in MF (*10–15*).

Since the transcriptome data set included in the MOFA model was restricted to 5000 genes (see methods section), we screened the entire transcriptomic data set to investigate signals of aberrant thymopoiesis in more detail. It has been reported that ectopic expression of *RUNX2* strongly expands immature thymocytes during the double negative stages, resulting in a preneoplastic state of thymocytes characterized by low proliferation rates (*62*). However, concomitant overexpression of *MYC* rescues proliferation and facilitates differentiation. Additionally, *MYC* and *RUNX* collaboratively inhibit the tumour suppressor p53, resulting in decreased apoptosis of malignant T cells. Together, this ultimately leads to the accumulation of mature, neoplastic T cells (*62*, *85*). In agreement with these reports, several genes were aberrantly expressed in the MF transcriptome (suppl. Material 1): Besides *RUNX1* and *RUNX2*, we identified enriched *MYCBP2* (Fig. 1D), which is a member of the c-myc family with the function to increase c-myc activity (*86*). Further, although the pathway to regulate p53 activity was upregulated (Fig. 1B), important components of the p53 machinery were downregulated (Fig. 1D): These included the p53 gene *TP53* itself, the p53 stabilizing protein *NOP53* (*87*), and TP53AIP1 which is regulated by p53 and induces apoptosis (*88*).

In summary, our data shows dysregulation of pathways involved in early thymopoiesis, probably representing the initial oncologic transformation of T cells in MF. Moreover, factor 5 suggested a connection between viruses and MF aetiology given the concomitant upregulation of host responses to viruses and *RUNX1/2* signalling.

### Increased viral prevalence and EBV-epitope recognition in MF skin lesions

In MF, malignant T cells circulate in the blood and infiltrate the upper dermis but have not been described to be present in the most superficial layers of the skin (*89*). To determine whether viruses may be involved in in MF aetiology, we investigated viral prevalence in different layers of the skin and the blood of MF patients. Viruses in superficial skin layers were identified through whole metagenomic sequencing (WMS) initially generated during our previous study (*41*). Viruses in deeper skin layers like epidermis and dermis as well as in the blood were identified thorugh RNAseq generated the present study.

In whole skin samples, total viral load trended higher in MF lesions compared to nonlesional skin (Fig. 4B), whereas no difference was observed in superficial skin samples (Fig. 4A). Among all viruses detected in MF lesions of whole skin samples, EBV and human papillomavirus 71 (HPV71) displayed the highest prevalence, albeit HPV71 was also frequently present in nonlesional skin. Remarkably, EBV and HPV71 were also present in the blood of some patients (Fig. 4B). Both, EBV and various human papillomaviruses are implicated in the development of cancer, including T cell lymphomas (*90–92*). Since MF is characterized by the infiltration of circulating T cells from the blood into the skin (*89*), the concomitant presence of viruses in both compartments indicates a potential viral involvement in MF.

**Figure 4:**
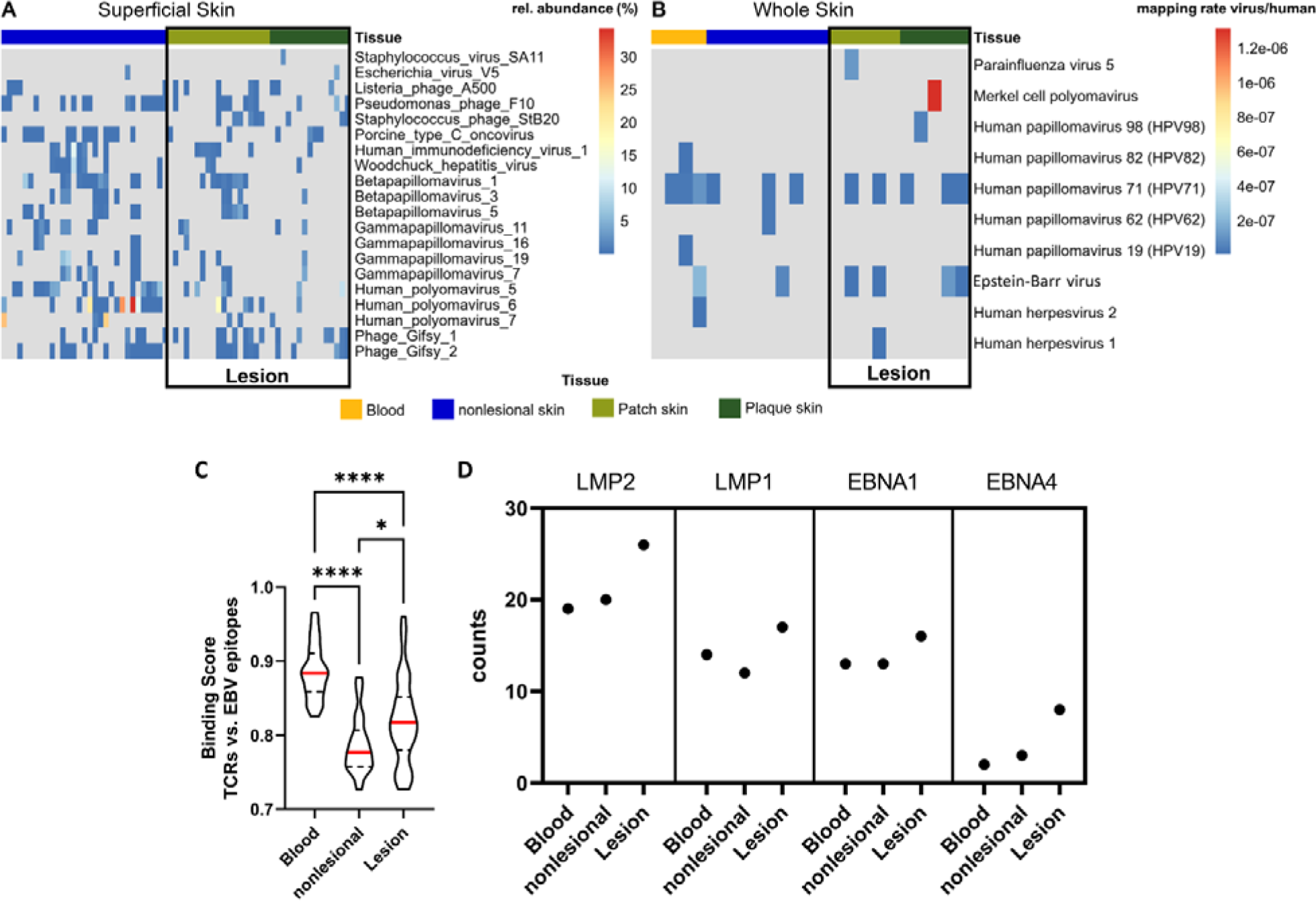
Viral association with MF. **(A – B)**, Viral prevalence identified in WMS reads of superficial skin **(A)** and RNAseq reads of whole skin **(B)**. Viral prevalence in MF lesions of whole skin trended higher compared to nonlesional skin, specifically EBV and HPV71 **(B)**, whereas no difference was present on superficial skin **(A)**. In **(A)**, the top 20 viruses with the highest variation between MF lesions and nonlesional skin are shown. In **(B)**, all viruses detected are shown. **(C – D)**, T cells in MF lesions and the blood specifically recognize EBV epitopes. **(C)** The probability of TCRs in blood, nonlesional skin and MF lesions to recognize EBV epitopes was assessed using ERGO-II (*94*). Binding Scores (1 = perfect binding, 0 = no binding) were calculated for each TCR with each EBV epitope. n = 175, displayed are violin plots with the median (red), 1^st^ and 3^rd^ quartile, Kruskal-Wallis test. **(D)** Displayed are the EBV-epitopes that were recognized most often by TCRs of MF patients.

Next, we evaluated whether T cells in deeper skin layers or T cells in the blood were directed against known EBV epitopes obtained from the Immune Epitope Database (IEDB) (*93*). We sequenced the TCR of MF lesions, nonlesional skin and blood and utilized ERGO-II (*94*) to calculate binding scores for the most abundant TCRs of a given sample with each of the EBV-epitopes. TCRs in MF lesions and the blood showed a significantly higher probability to recognize EBV-epitopes than T cells in nonlesional skin (Fig. 4C), which agreed to the increased viral prevalence in deeper layers of MF lesions. Interestingly, latent membrane protein (LMP) 1, LMP2, and Epstein-Barr-nuclear antigen (EBNA) 1 were most frequently recognized EBV-epitopes by TCRs in the skin and blood (Fig. 4D). The expression pattern of LMP1, LMP2, and EBNA1 represents a characteristic EBV gene profile of latency type I/II, commonly observed in EBV-induced T cell lymphomas (*95*, *96*). In NK/T Cell lymphoma, LMP1 upregulates *CD274* (PD-L1) via the NF-κB axis (*97*), which agreed with our results of enriched *CD274* and NF-κB signalling in plaque stage (Fig. 1B, D).

Together, the trend of increased EBV and HPV71 prevalence in deeper skin layers along with a significantly higher probability of EBV-epitope recognition by TCRs in MF lesions and the blood compared to nonlesional skin suggest viral involvement in MF aetiology.

## Discussion

The clinical course of MF varies greatly, with some patients having only minor progress, whereas others suffer from fast progression and high disease burden (*5*, *98*). Likewise, the transcriptome is tremendously heterogeneous, exhibiting patient and even lesion specificity (*27*, *39*, *53*). Moreover, ambiguity exits about the initial oncologic transformation of malignant T cells, as studies from different groups mapped the event to either early thymocytes (*10–15*) or mature, effector memory T cells (*9*). Consequently, a common pathogenic mechanism of all patients or patient subgroups is still in doubt, resulting in a non-optimal treatment regimen (*27*). We recently demonstrated that the skin microbiome of MF patients is altered and identified a subgroup of patients that was overgrown by a distinct, pathogenic *S. aureus* strain. This subgroup exhibited a significantly aggravated disease course, possibly owing to the virulence factor spa which was present in the *S. aureus* genome (*41*). In line, others showed that spa can activate the NF-κB axis (*42*), which is recurrently deregulated in MF patients with aggressive disease (*34*, *44–46*, *99*). We thus theorized that (I) the lesion-specific microbiome determines transcriptomic response, thereby contributing to or causing heterogeneity (*40*) and that (II) spa-bearing *S. aureus* elicits NF-κB signalling to fuel MF. Therefore, we investigated how the skin microbiome affects the skin transcriptome in a subset of 10 MF patients that were enrolled in our previous study (*41*). Using RNAseq, multiomic data integration, virome profiling and TCRseq, we obtained novel insights into the role of the microbiome in both the aetiology and pathogenesis of MF.

First, we recovered substantial transcriptomic heterogeneity on both gene- and pathway-level and observed that this heterogeneity may be largely driven by differential expression of T cell signalling pathways. Strikingly, our results show this differential T cell signalling pattern was caused by the skin microbiome, since denoising the transcriptome with the microbiome using MOFA strikingly reduced heterogeneity (Fig. 2A, B)

Second, latent factor 4 strongly suggested that *S. aureus* elicited the upregulation of both non-canonical NF-κB and IL-1B signalling, which explains the aggravated disease course of MF patients overgrown by a spa-bearing *S. aureus* strain. Non-canonical NF-κB signalling is known to promote the survival and proliferation of thymocytes and mature T cells, induce the differentiation of naïve T cells into effector memory T cells, and support clonal expansion (*69*, *70*), which are all common characteristics of malignant T cells in MF (*1*, *8*, *9*, *100*). NF-κB signalling is typically initiated by the interaction of TNFα with either TNFRSF1A (also known as TNFR1) or TNFRSF1B (also known as TNFR2) (*69*, *70*). However, the enrichment of non-canonical NF-κB signalling in factor 4 appeared to be independent of TNFα, since it displayed a very low weight. Instead, we propose that the *S. aureus* virulence factor spa induced non-canonical NF-κB signalling since Gómez et al. showed that spa activates TNFRSF1A (*42*, *43*). In general, TNFRSF1A induces the canonical form of NF-κB signalling leading to apoptosis, while TNFRSF1B induces non-canonical NF-κB signalling resulting in survival and proliferation. However, there is some level of crosstalk between the two forms of NF-κB pathways. In highly activated T cells, such as the malignant T cells in MF, activation of TNFRSF1A results in survival rather than apoptosis (*69*, *70*, *101*). Further, the extracellular domains of TNFRSF1A and TNFRSF1B exhibit a high degree of structural similarity (*102*), possibly allowing spa to activate both receptors. Additionally, while TNFRSF1A is expressed nearly ubiquitously across various cell types throughout the body, TNFRSF1B expression is considerably more restricted, including thymocytes and T cells (*70*). Thus, spa may activate either TNFRSF1A, TNFRSF1B or both to initiate non-canonical NF-κB signalling to fuel MF progression.

In agreement with our findings, a clinical study of CTCL patients reported that systemic inhibition of NF-κB induced skin response in 30.4% of patients whereas blood response was mixed (*103–105*) Further, Shin et al. identified ectopic NF-κB signalling in 30.6% of MF patients and that these patients had an aggravated disease course compared to patients without ectopic NF-κB signalling (*44*). We previously identified that 31.3% of MF patients were overgrown by the virulent, spa-bearing *S. aureus* strain and had an aggravated disease course (*41*). Given the similar prevalences of the described subgroups and the likely NF-κB activating effect of spa indicates a possible connection between skin colonization by a spa-bearing *S. aureus* strain and disease aggravation in the aforementioned studies. Additional research on spa-NF-κB interaction including mechanistic assays with bigger cohorts is warranted.

Regarding the upregulated IL-1B signalling in factor 4, It was shown that *S. aureus* induces secretion of IL-1B by eosinophiles (*106*), which infiltrate MF lesions (*107*). Furthermore, human keratinocytes, skin derived dendritic cells and lymphocytes produce several cytokines including IL-1A, IL-1B, and IL-4 when challenged with *S. aureus* or spa (*71*, *108*, *109*). This suggests a paracrine stimulation of malignant T cells by the tumour microenvironment via cytokine signalling in response to *S. aureus*.

Last, we discovered evidence indicating viral involvement in the aetiology of MF. Latent factor 5 captured a concomitant upregulation of innate anti-virus defence mechanisms and aberrant *RUNX1/2* signalling. The latter is known to coordinate thymopoiesis during double negative (DN) stages DN1 through DN3 (*62*, *63*, *110*), which have been mapped to the initial oncologic transformation in MF (*10–15*). We further observed that viral load, particularly of HPV71 and EBV, trended higher in both deeper skin layers of MF lesions and the blood compared to nonlesional skin, whereas superficial skin layers of MF lesions showed no difference. HPV71 was shown to degrade p53 (*111*), which can result in neoplasia (*112*). Nevertheless, while certain papillomaviruses are categorized as high-risk factors for the onset of solid cancers (*113*) and have also been loosely linked to an elevated risk of lymphomas (*114*), HPV71 is generally regarded as having low oncogenic potential (*115*). In contrast, EBV is a well-known oncovirus (*116–118*). Remarkably, we found that T cells residing in MF lesions or in the blood were significantly more affinitive to EBV-epitopes than T cells of nonlesional skin (Fig. 4C). Moreover, the most frequently recognized EBV-epitopes were epitopes of LMP1, LMP2, and EBNA1 which are typical genes expressed by EBV in latent infection (*96*). Although EBV has a strong B cell tropism, leadting to B cell Hodgkin’s lymphoma (HL) and non-Hodgkin’s lymphoma (nHL) (*116–118*), the virus also infects T cells thereby causing some entities of T cell nHL (*47*, *48*). Notably, CTCL patients have an increased risk of developing secondary HL and nHL (*17–20*). It has been proposed that a single precursor malignant T cell can trigger HL, CTCL and lymphomatoid papulosis, a benign T cell neoplasm, in the same patient (*21*, *22*). Others reported the simultaneous presence of HL and CTCL within the same lymph node and theorized that the two malignancies arise from distinct B and T cells (*23*), indicating that a common trigger induced oncogenesis.

To the best of our knowledge, we were the first to investigate viral prevalence in different skin compartments and found that EBV and HPV71 trended higher solely in deeper skin layers of MF lesions, where malignant T cells in MF reside (*89*). Collectively, our data suggests that viruses, probably EBV and/or HPV71, play a role in MF aetiology. Further investigations with larger cohorts and longitudinal viral monitoring across both tissues and skin compartments are necessary to determine whether viruses are the aetiologic agent in MF.

## Conclusion

In this study, we provide several lines of evidence that the skin microbiome influences MF. First, our results demonstrated that the skin microbiome largely contributes to transcriptional heterogeneity. Second, we showed that a spa-bearing *S. aureus* strain, which overgrows a subgroup of MF patients with aggravated disease course, evokes non-canonical NF-κB and IL-1B signalling in the skin. Third, our data collectively indicated that viruses, particularly EBV and HPV71, may be the aetiologic agent in MF. Together with results from our previous study, these findings led us to propose a model of microbiome-driven MF aetiopathogenesis (Fig. 5): The initial oncologic transformation emerges during early thymopoiesis triggered by aberrant *RUNX* expression, potentially caused by viral infection such as EBV and/or HPV71. Malignant T cells infiltrate the skin, leading the microenvironment to release AMPs. Subsequently, this process eliminates microbes, resulting in skin dysbiosis and a diminished epithelial barrier. Additionally, certain AMPs attract CD4+ T cells, including potentially malignant T cells, thereby augmenting the infiltration of (malignant) T cells into MF lesions. Over time, some microbes acquire resistance to AMPs and recolonize the lesions. In the ΔSA-neutral subgroup, microbes with anti-*S. aureus* properties accumulate, resulting in a balanced microbiome and favourable clinical course. However, in the ΔSA-positive subgroup, a virulent *S. aureus* strain carrying the virulence factor spa outgrows and activates non-canonical NF-κB and IL-1B signalling, resulting in the generation of mature effector memory T cells and poor outcome. Hence, both perspectives of the cells of origin in MF may apply, early T cell progenitors (*10–14*) and mature, effector memory T cells (*8*, *9*). Limitations of our study are the small patient cohort, the mono-centric nature of our study, and the lack of mechanistic assays to confirm our findings which are based on computational analyses from primary clinical specimens. Further research is needed to understand how the microbiome influences or contributes to MF aetiopathogenesis.

**Figure 5:**
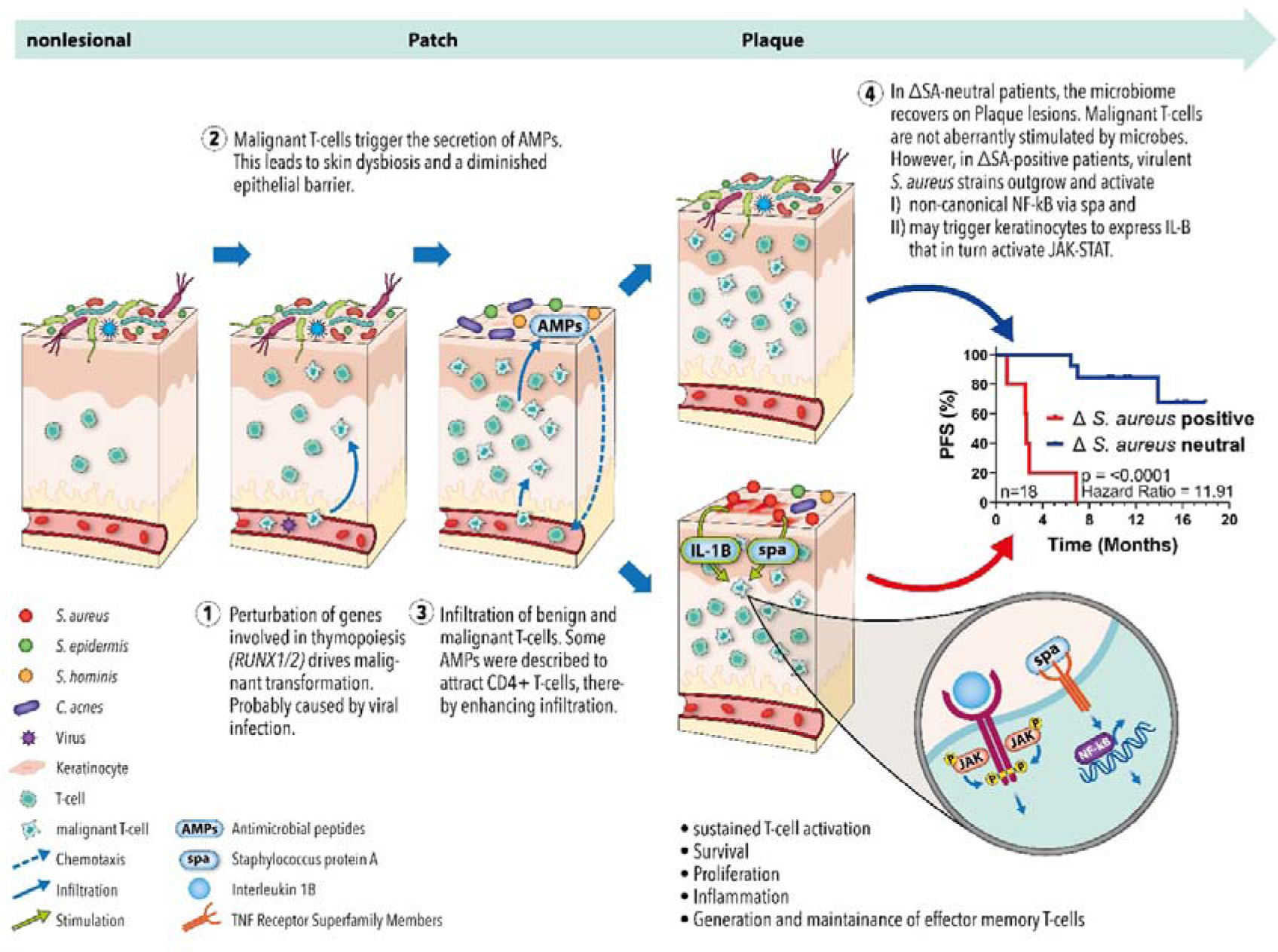
Proposed model of microbiome-driven MF aetiopathogenesis. T cell precursors undergo initial oncologic transformation due to aberrant expression of *RUNX1* and *RUNX2*, probably caused by viruses like EBV and/or HPV71. After maturation and skin infiltration, the malignant T cells trigger the microenvironment to secrete AMPs. AMPs kill most of the physiological skin microbiota, resulting in dysbiosis and a diminished epithelial barrier. In addition, AMPs may recruit benign and malignant CD4+ T cells, establishing a loop of sustained dysbiosis and T cell infiltration. Since AMP levels remain constant over the disease course, some microbes eventually acquire resistance and recolonize the lesions. In the ΔSA-neutral subgroup, microbes with anti*-S. aureus* activity accumulate, resulting in a balanced microbiome that does not fuel the disease. However, in the ΔSA-positive subgroup, virulent *S. aureus* strains bearing the virulence factor spa overgrow and activate non-canonical NF-κB as well as IL-1B signalling. The two pathways cause inflammation, sustained activity, survival, and proliferation of T cells, as well as the generation and maintenance of effector memory T cells. These characteristics are hallmarks of malignant T cells in MF and may explain the significantly aggravated clinical course of the ΔSA-positive subgroup.

## Methods

### Study Design and Clinical specimens

The study was designed as case-control study using intra-patient controls to account for the high variability of microbiome profiles between patients (*41*). The patient cohort comprised a subset of patients included in our previous study (*41*) that were recruited from the Department of Dermatology, University Medical Centre Mainz, Germany. Metagenomic samples were obtained with a swab-scrape-swab procedure described earlier (*41*). In brief, a pre-moistured swab was brushed over the skin, then the same skin area was gently scarped with a scalpel, and then brushed again with the same swab. 4 mm punch biopsies were obtained from the same MF lesions sampled for the metagenome. Peripheral blood was drawn for the isolation of peripheral blood mononuclear cells (PBMCs) which were subsequently enriched for the T cell fraction detailed in reference (*41*).

All procedures in this study were conducted in accordance with the Declaration of Helsinki from 1964 and its later amendments. The local ethics committee, the Medical Association of Rhineland-Palatine (Ethik-Kommisssion der Landesaerztekammer Rheinland-Pfalz), Germany, assigned ethical approval under number 2020-14813.

### RNA sequencing and Transcriptome Analysis

Total RNA was extracted from skin biopsies and the enriched T cell fraction as described previously (*41*). Total RNA was sent to Novogene Company Ltd. (United Kingdom) for library preparation and sequencing. Briefly, after quality check with Caliper Life Sciences GX II (USA), 400 ng RNA were used as input for NEBNext Ultra RNA Library Prep Kit (New England Biolabs, USA). One sample (nonlesional skin of Pat1) was excluded due to low RNA quality. After quality check libraries were sequenced on a NovaSeq 6000 (Illumina, USA) at 150 bp paired-end to an average of 78.74 (range 63.82 – 94.20) million raw reads, 11.81 (range 9.57 – 14.13) giga base pairs (Gbp) per sample.

For the transcriptome analysis, raw reads were quality checked with Fastqc version 0.11.9 (*119*), and subsequently aligned against the human reference genome Ensembl built GRCh38.p13 using STAR 2.7.9a with the flag –quantMode GeneCounts (*120*). The data was loaded into DESeq2 version 1.36.0 (*121*), filtered for genes with a minimum of ten cumulative counts over all samples, and normalized using the internal DESeq2 method. Subsequently, a differential expression analysis using the Wald test was carried out, adjusted for influences from individual patients which may have been introduced by the paired-sample design. Genes with <0.05 adjusted p-value were considered significant. Volcano plots were created using the R package EnhancedVolcano (*122*). For principal component analyses, the data was first transformed with the variance-stabilization method and then analysed using the plotPCA function within DESeq2. The PCA was visualized using ggplot2 (*123*).

Gene set enrichment analysis was performed using the R package pathfindeR (*124*). Log2 fold-changes of significantly deregulated genes determined by DESeq2 were used as input. The Reactome database was used to find enriched pathways (*125*). Because we were particularly interested in signalling pathways that might orchestrate MF pathogenesis, we screened the literature for pathways that were described as deregulated in T cell lymphomas or CTCL. The Reactome database was subsequently filtered to retain only pathways that contain at least one of the following terms (case insensitive): cascade, signal, signalling, signaling, pathway, transition, cycle, regulation, activation, keratinization, cornified, antimicrobial, interferon, IFN, stimulation, stimuli, activate, receptor, TLR. Enriched pathways were clustered using the pathfindeR function cluster_enriched_terms. Plots were created using pathfindeR.

### Multiomic Data Integration

Two data modalities, the microbiome and the transcriptome, were integratively analysed using the R implementation of MOFA version 1.6.0 (*61*). The microbiome dataset was created in our previous study where we characterized the MF skin microbiome using shotgun metagenomics and MetaPhlAn 3.0.2 (*41*, *126*, *127*). To obtain absolute counts of microbial taxa (rather than the relative abundance, which is the default in MetaPhlAn and was used in our previous study), the metagenome was re-profiled including the flag -t rel_ab_w_read_stats using MetaPhlAn 3.0.2 with the intermediate bowtie2 mapping files as input. The resulting profiles were filtered to retain only taxa on the species level and normalized using the wrench method, which accounts for sparse metagenomic count data and is implemented in the R package wrench (*128*). The normalized data was log2 transformed with a pseudocount of 1. To select highly variable species for the MOFA model, 50 species with the highest variance were filtered (with the R base function var) and used for all subsequent steps. The transcriptome data set was created in this study as described above. After mapping and normalization with STAR 2.7.9a and DESeq2 version 1.36.0 (described above), the data was transformed with the regularized logarithm using the rlog function implemented in DESeq2 (*121*). As proposed by the MOFA authors, the large transcriptome data set was restricted to not overrepresent the smaller microbiome data set in the MOFA model. Therefore, the top 5000 highly variable genes were selected based on the median absolute deviation calculated with the R base function mad.

The MOFA model was trained with a single group, the option scale_views turned true, convergence_mode slow, and eight factors. Gene set enrichment analysis was performed with the MOFA function run_enrichment, which is based on principal component gene set enrichment (PCGSE) (*129*) using only genes with positive weights as input. The Reactome database filtered for pathways of interest (see above section of transcriptome analysis) was used as background. Pathway enrichment plots were generated with ggplot2 and all heatmaps were created with the R package pheatmap.

### Virus Profiling

The virome present on superficial skin layers was profiled via re-analysis of the microbiome dataset generated in our previous investigation (*41*). Briefly, human skin was brushed topically with swabs, and whole metagenomic sequencing was performed and subsequently profiled with MetaPhlAn 3.0.2. While viruses were excluded in our previous study, here we activated the flag for viral identification (--add_viruses). The resulting profiles were filtered to retain only viruses.

The virome present in deeper skin layers like the dermis and epidermis was profiled using RNAseq reads from skin punch biopsies obtained in the present study. To this end, the pipeline VIRTUS was applied (*130*). Briefly, non-human reads were filtered out via mapping RNAseq reads to the human reference genome Ensembl built GRCh38.p13. Next, unmapped (i.e., non-human) reads were aligned against 762 viral genomes. The heatmaps were generated with the R package pheatmap (*131*).

### Assessment of TCR – Epitope Binding

To investigate whether T cells of MF patients recognize EBV-epitopes, the T cell receptor was sequenced as described previously (*41*). Briefly, RNA isolated from skin and blood of MF patients were subjected to library preparation spanning the variable part of the TCR using the NEBNext Immune Sequencing Kit, human (E6320, New England Biolabs, USA) and sequenced on a MiSeq (Illumina, USA) running 300 bp PE. The sequencing data was processed with the pRESTO toolkit (*132*) (https://usegalaxy.org/u/bradlanghorst/w/presto-nebnext-immune-seq-workflow-v320) and the R package immunearch (*133*). EBV-epitopes were obtained from the Immune Epitope Database (IEDB) (https://www.iedb.org/) (*93*). The Tool ERGO-II (*94*) was used to determine binding scores of pairs of TCRs and EBV-epitopes. The closer the binding score is to 1, the higher the probability that a TCR recognizes the epitope (*94*).

## Data Availability

All data modalities used in this study, RNAseq, WGS, and TCRseq, were deposited on the Gene Expression Omnibus (GEO) under the SuperSeries GSE221150 (https://www.ncbi.nlm.nih.gov/geo/query/acc.cgi?acc=GSE221150).

RNA sequencing data and associated analysis files can be accessed under GSE221148 (https://www.ncbi.nlm.nih.gov/geo/query/acc.cgi?acc=GSE221148). This includes viral profiles from both whole skin biopsies and the blood.

Microbiome sequencing data and associated analysis files can be accessed under GSE221149 (https://www.ncbi.nlm.nih.gov/geo/query/acc.cgi?acc=GSE221149). This includes re-analysed microbiome profiles with absolute counts and viral profiles from superficial skin layers.

TCR Sequencing data and associated analysis files can be accessed under GSE218874 (https://www.ncbi.nlm.nih.gov/geo/query/acc.cgi?acc=GSE218874).

## Acknowledgments

We thank Stefan Kindel, University Medical Centre Mainz, for the creation of the figure summarizing our proposed model of microbial-driven MF aetiopathogenesis.

