## Supplementary material for "Multiomic Data Integration Reveals Microbial Drivers of Aetiopathogenesis in Mycosis Fungoides": Suppl. Material 4

### Fitting a Multi-Omic Factor Analysis (MOFA) model to disentangle the influence of the microbiome on the transcriptome

To assess the influence of the microbiome on the transcriptome, we employed Multi-Omic Factor Analysis (MOFA). The MOFA model explained ~70% of the total variation in the transcriptome and ~30% of the total variation in the microbiome (suppl. Figure 1). Furthermore, the model recovered patterns in the multi-omic data set which we also observed in the single analyses of the transcriptome and the microbiome (*1*), showing a good fit of the model.

In particular, **Factor 1** likely matched MF pathomechanisms that are not dependent on microbial stimuli since it captured the majority of the variation in transcriptome but only little variation in the microbiome(Fig. 3A). MF archetypical genes like *CXCR4* (*2*) displayed high weights (suppl. Material 5) and GSEA showed several enriched pathways that we identified in plaque stage, like TCR signalling, CD28 co-stimulation, cytokine signalling, and PD-1 signalling (Fig. 3C). **Factor 2** resembled the signatures of patch stage, as it was characterized by solely negative or neutral microbiome feature weights (Fig. 3B), corresponding to the depletion of majority of microbes on patch lesions (*1*). Further, factor 2 was defined by enriched pathways involved in building the cutaneous barrier as well as TCR signalling and Interferon signalling (suppl. Material 5), agreeing to the transcriptome analysis of patch stage (Fig. 1B, supp. Material 2). **Factor 3** and **factor 4** resembled the microbiome patterns on plaque lesions of the ΔSA-neutral and ΔSA-positive subgroup, respectively. In factor 3, *S. aureus* displayed a negative weight while species with anti-*S. aureus* activity like *S. hominis, C. acnes* and *C. amycolatum* displayed positive weights (*3*–*7*), which was vice versa in factor 4. In our previous study we identified an aggravated disease course in the ΔSA-positive subgroup likely owing to the outgrowth of *S. aureus*, whereas in the ΔSA-neutral subgroup *S. aureus* abundance was stable. In line, the transcriptome of factor 3 showed no signatures of MF disease progression, but rather genes involved in skin development or scar/fibrous tissue formation (*8*, *9*) (suppl. Material 5), underpinning the favourable clinical course of the ΔSA-neutral subgroup (*1*).

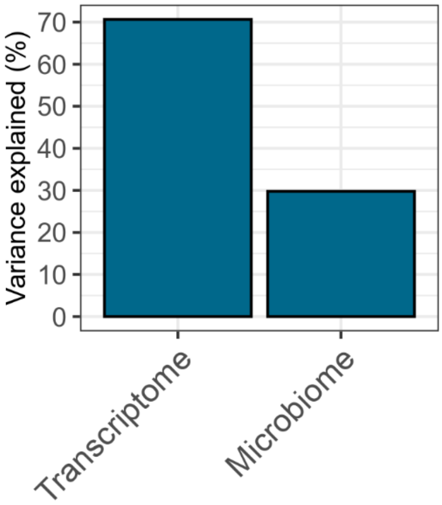


Suppl. Figure 1: Total variance explained in the transcriptome and the microbiome data modalities by the MOFA model.
