## Supplementary figures and images for "Multiomic Data Integration Reveals Microbial Drivers of Aetiopathogenesis in Mycosis Fungoides"

### Patch_GSEA_EnrichmentComparison.pdf

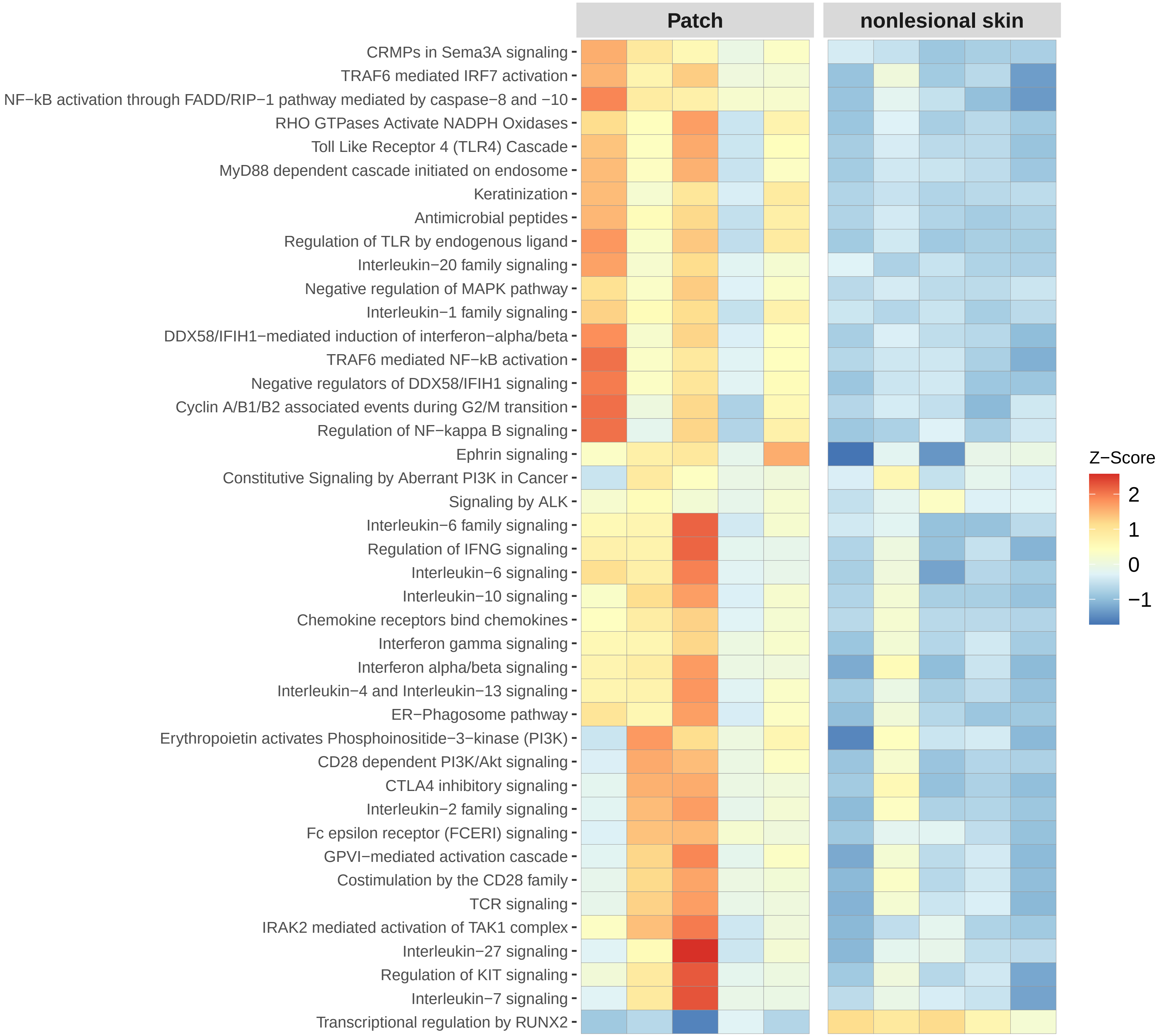

### Plaque_GSEA_EnrichmentComparison.pdf.pdf

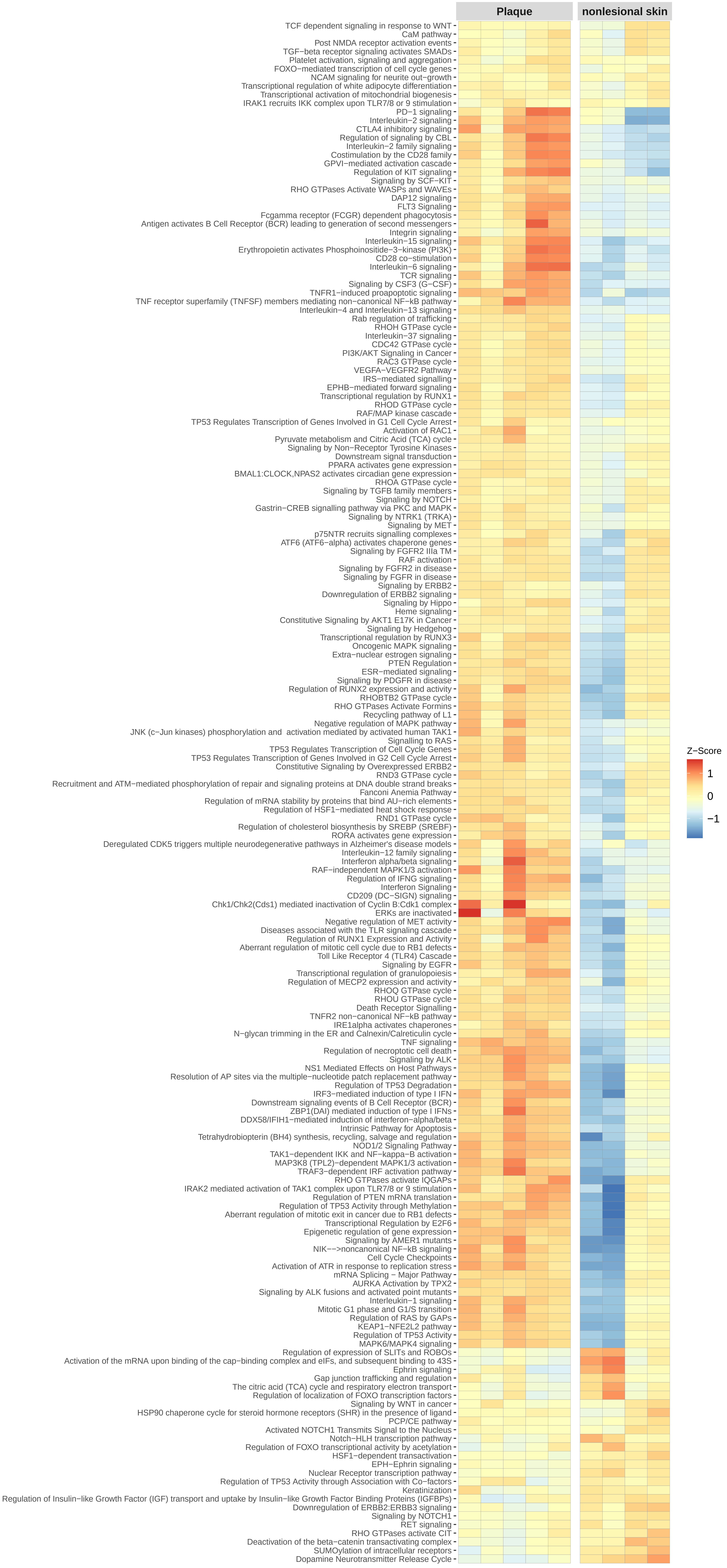
