## Supplementary material for "Multiomic Data Integration Reveals Microbial Drivers of Aetiopathogenesis in Mycosis Fungoides": Suppl. Material 5: Suppl_Material5_Plots.pdf

#### Enriched Pathways and genes with the highest weight in each pathway of Factor 1

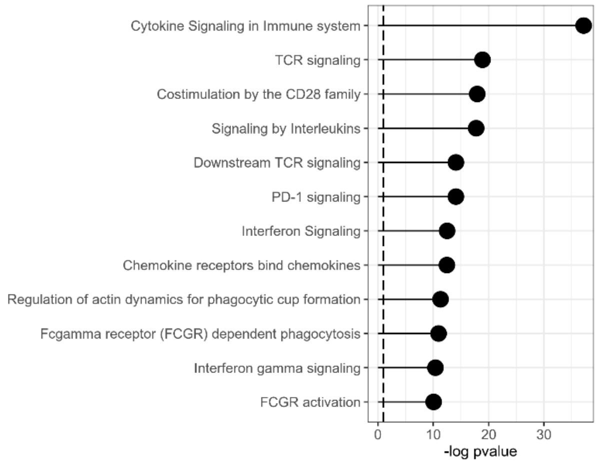


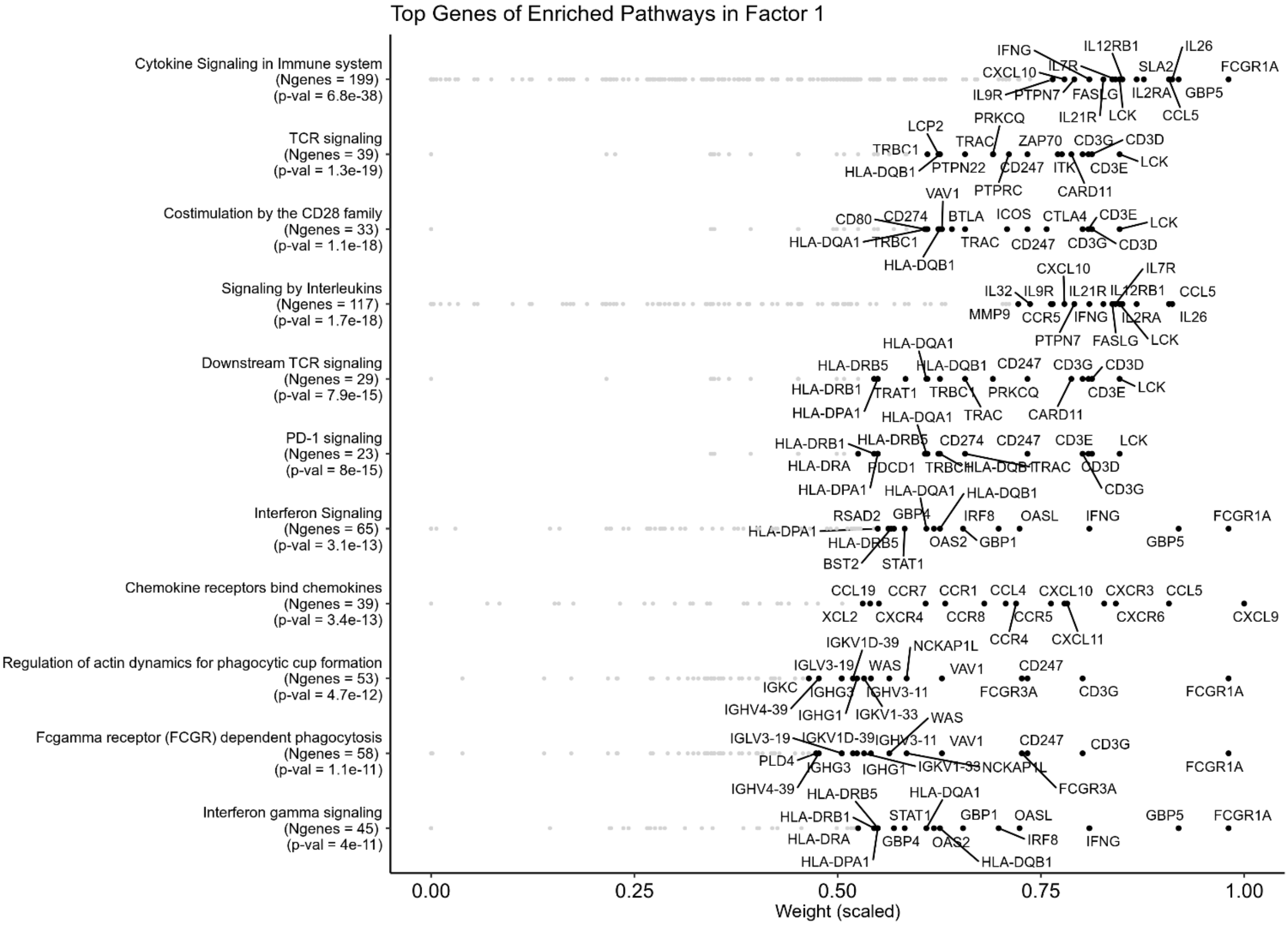

#### Enriched Pathways and genes with the highest weight in each pathway of Factor 2

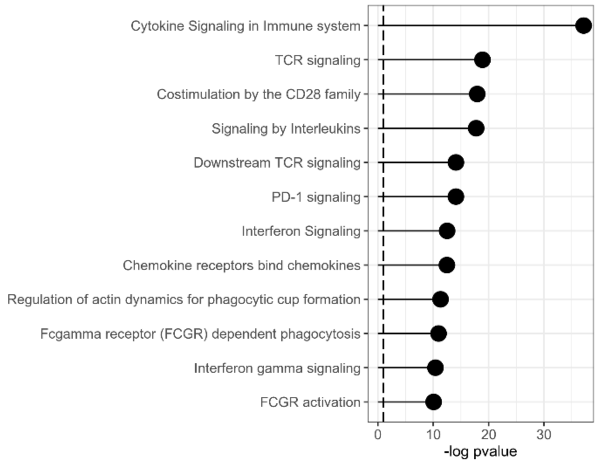


Top Genes of Enriched Pathways in Factor 2

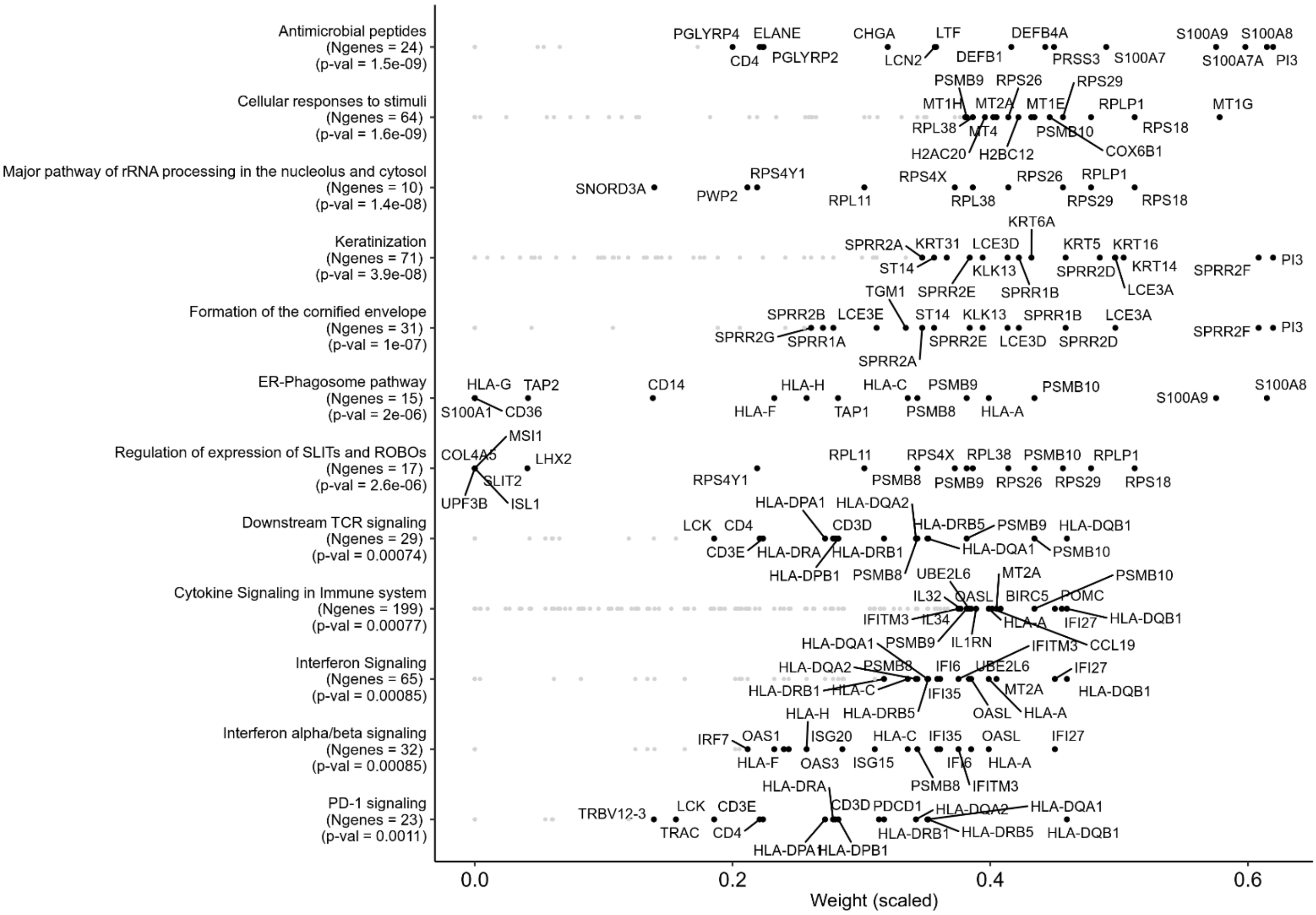

#### Enriched Pathways and genes with the highest weight in each pathway of Factor 3

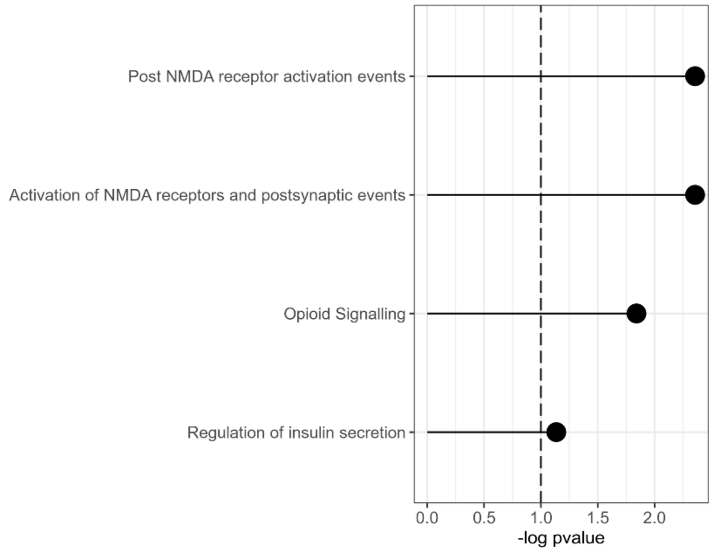


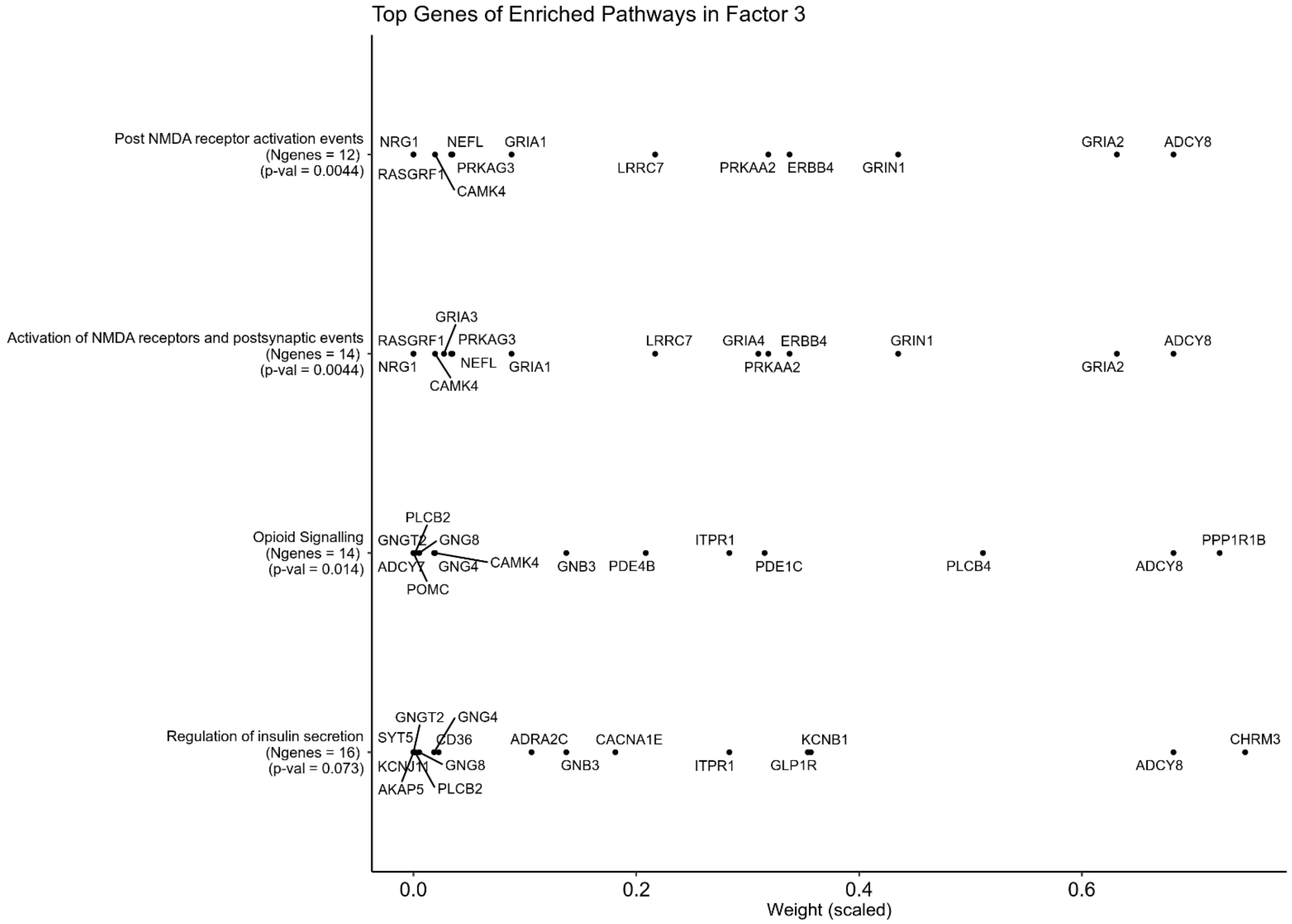

#### Enriched Pathways and genes with the highest weight in each pathway of Factor 4

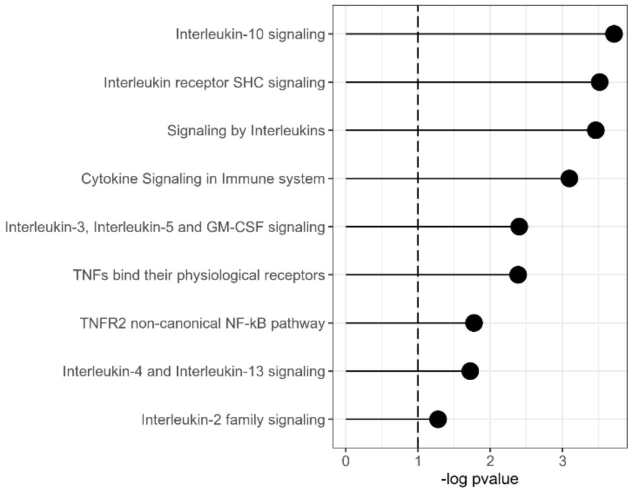


Top Genes of Enriched Pathways in Factor 4

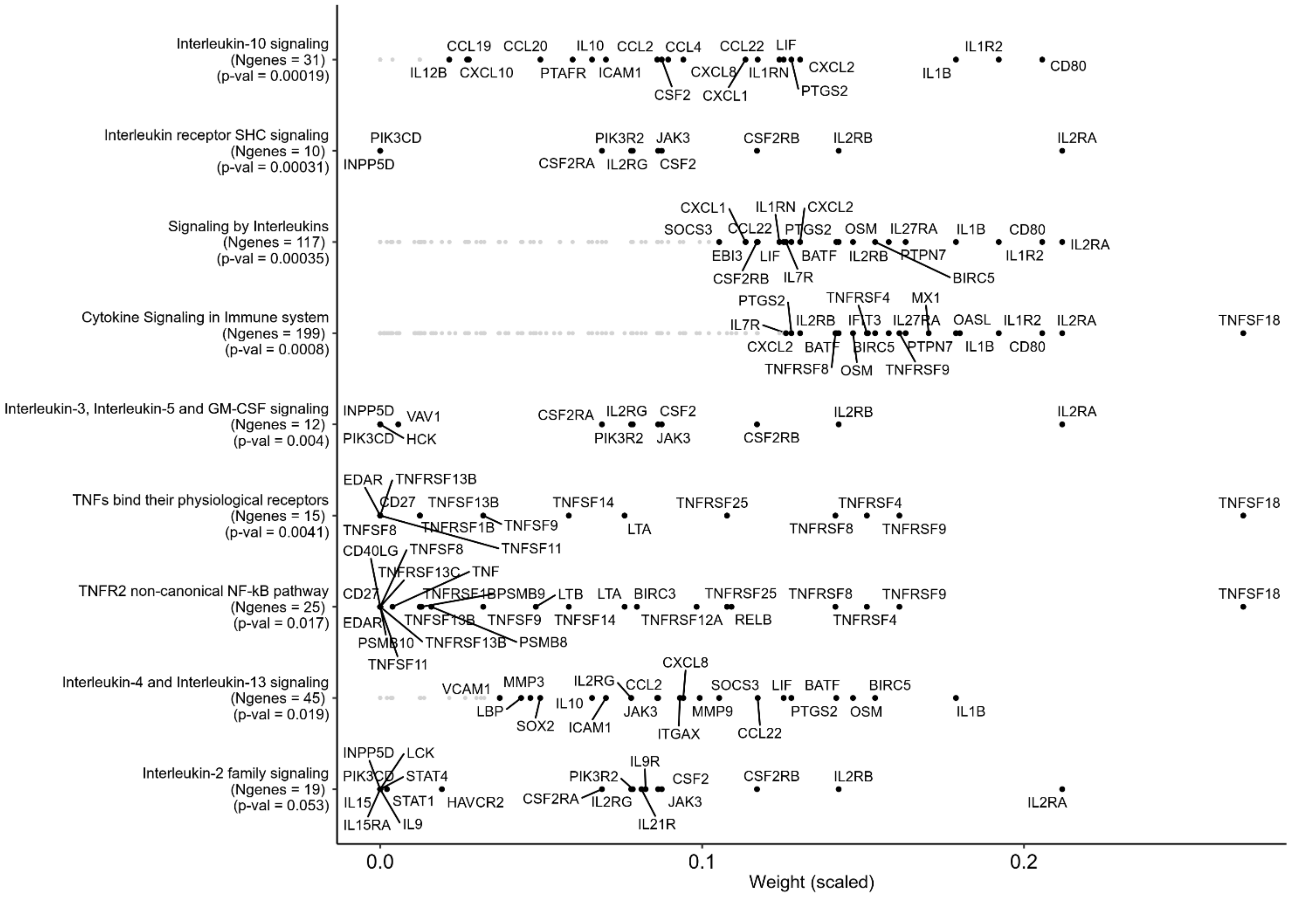

### Enriched Pathways and genes with the highest weight in each pathway of Factor 5

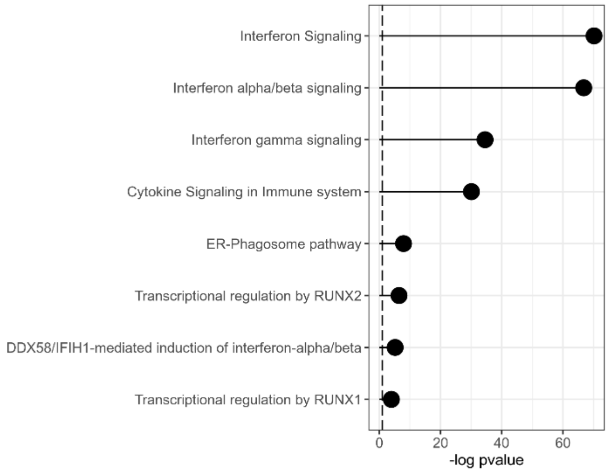


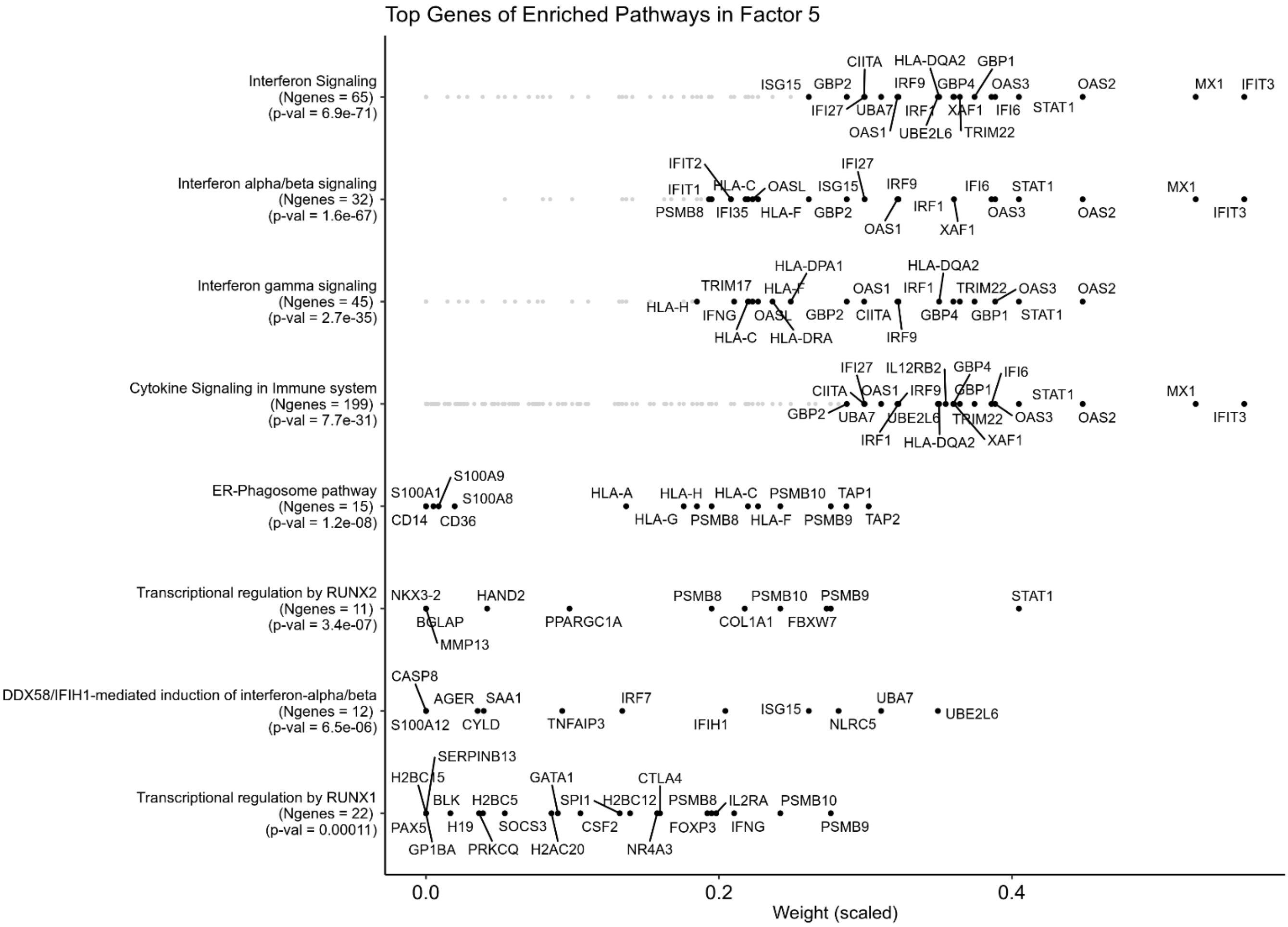
